# Effect of cellular nutrient economy on the evolution of genome size in phytoplankton

**DOI:** 10.1101/2025.09.15.676320

**Authors:** Carlos Caceres, Marc Krasovec, Olivier Crispi, Sebastien Gourbiere, Gwenael Piganeau

## Abstract

The origin of genome size variation remains a central question in evolutionary biology. Previous analyses suggest that the energetic costs of small insertions and deletions (indels) influence genome size by altering the cellular energy budget. Here, we present a theoretical framework for genome evolution in phytoplankton—the main producers of the sunlit ocean—whose growth depends on nutrient availability. We derive an expression for the selection coefficient on indels as a function of the minimum amount of nutrient inside the cell. Parameter estimates indicate that 1-bp indels can be counterselected in phytoplankton with minimal nutrient requirements. We then test three predictions of nutrient-driven selection on genome evolution. Altogether, this model provides a quantitative and mathematical basis for the genome streamlining hypothesis.

## Introduction

Genome size spans over five orders of magnitude, from 1.3 Mb in prokaryotes to more than 100 Gb in certain eukaryotes ^1,2^. Such a variation is primarily driven by differences in the size of non-coding regions ^3–5^. Present genome sizes result from past genome expansion and compaction dynamics driven by molecular processes generating “indel” mutations and evolutionary processes shaping their fixation ^3^. Here, we use “indel” in a broad sense to refer to any mutation generating a genome size variation, which includes deletions, insertions, duplications, hybridizations or lateral gene transfers.

Many post-hoc explanations of the diversity of genome sizes have been suggested ^5^. Of relevance for microorganisms, the “streamlining hypothesis” states that small genome sizes may have evolved to minimize the material cost of replication, leading to the selection of species with smaller genomes in resource-limited environments ^1,6–9^. However, this hypothesis currently lacks a mechanistic theoretical framework linking genome size to the fixation probability of indels, which depends on both natural selection and genetic drift ^3,4^. This framework requires estimating the selection coefficients of indel mutations. Previous approximations were based on the relative energetic cost of indels compared with a cell”s ATP requirements from glucose ^10,11^. Here, we provide an alternative estimation based on a model linking the molecular changes in the number of nitrogen (N) and phosphate (P) atoms in an indel to differences in the growth rate between the mutant carrying the indel and the wild type. In this way, the selection coefficients of indels can be explicitly linked to the changes in cellular nutrient requirements they cause.

The recent increase in genome resources in phytoplankton allows renewed investigation into the role of cellular nutrient requirements on genome evolution. Phytoplankton are mostly unicellular and inhabit ecosystems that show a wide range of nutrient availability. Low N and P levels limit phytoplankton growth in vast areas of the ocean and drive genomic adaptations that enhance survival and resource utilization in nutrient-limited environments ^12–15^. Moreover, phytoplankton include some of the smallest genomes among free-living prokaryotes and eukaryotes: <2 Mb genomes in *Prochlorococcus* ^16^ and 13 Mb in *Ostreococcus* ^17,18^. Importantly, the growth rate of phytoplanktonic organisms can be described as a function of the intracellular nutrient content, or quota, of the most limiting nutrient by using the quota model ^19^. The model”s key parameter, *Q*_min_, is the minimum amount of nutrient inside the cell, required for population growth, and has been estimated for several phytoplankton species ^20^. Phytoplankton thus provide a unique opportunity to estimate selection coefficients (*s*) of indel mutations based on the changes in cellular nitrogen and phosphate content they cause.

### Non-coding regions drive between-species genome size variations in phytoplankton

To investigate genome size variations in phytoplankton, we analyzed 173 genomes from 28 bacteria and 101 eukaryotic species, spanning 5 of the 8 major photosynthetic lineages ^21^. Cyanobacterial genomes range from 1.6 to 8 Mb, whereas eukaryotic genomes span from 13 Mb in *Ostreococcus tauri* (Chlorophyta) to 4.1 Gb in *Prorocentrum cordatum* (Dinoflagellate) (Fig. 1A and fig. 1A). In eukaryotes, genome size variation is primarily driven by differences in non-coding DNA content, ranging from 2.2 Mb (17.6%) in *O. tauri* to >4 Gb (96.9%) in *P. cordatum* (Fig. 1A,B and fig. S1A). Both intergenic and intronic regions contribute substantially to genome size, averaging39% and 20% of eukaryotic genomes, respectively); in 80% of annotated species, intergenic DNA exceeds intronic DNA (39 of 49 fig. S1B). In fact, genome size correlates positively with the average size of both intergenic and intronic regions (Fig. 1C, D). Lineage-specific patterns are also evident. In diatoms, coding and non-coding DNA contribute nearly equally to genome length (49% and 51% on average, respectively), whereas chlorophytes with similar genome sizes have a higher fraction of non-coding DNA (62% and 38%; Fig. 1A, B and fig. S1A). However, diatom genomes have a higher proportion of intergenic regions than Chlorophyta with a similar genome size (ANCOVA, F_1,14_ = 7.41, p < 0.05; Fig. f1B), reflecting a higher gene number and fewer introns per gene (ANCOVA, genes: F_1,14_ = 8.98, p < 0.01, Fig. 1E; introns per gene: F_1,14_ = 30.17, p < 0.001). Taken together, genome sizes in unicellular phytoplankton span four orders of magnitude, variation that largely reflects differencesin non-coding DNA content. This pattern is consistent with previous datasets spanning both unicellular and multicellular organisms ^4,5,22–24^.

**Fig. 1.**
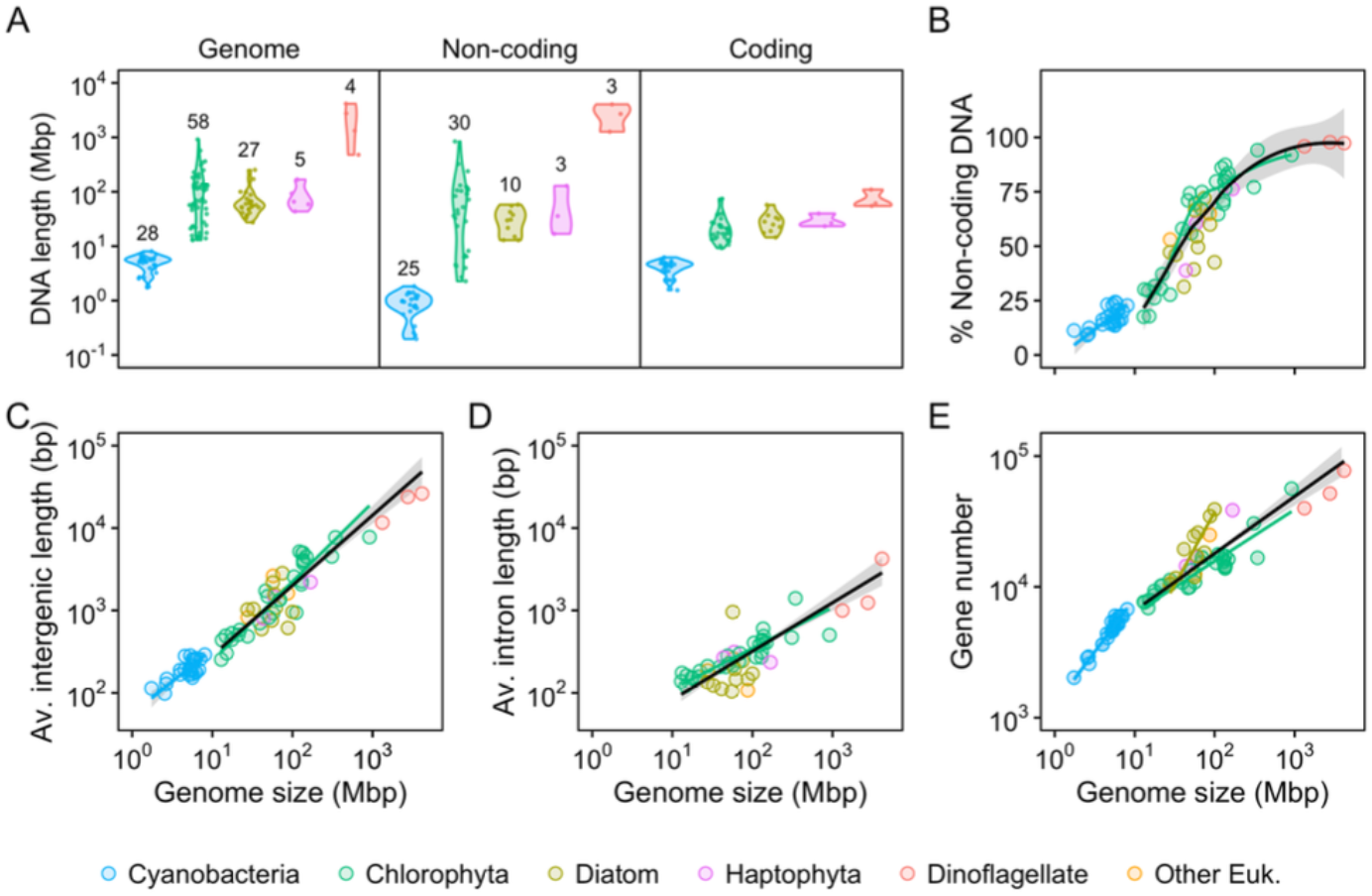
Size of DNA regions and their relationship with genome size. **(A)** Violin plots showing the size distribution of genomes, non-coding regions, and coding regions. Numbers indicate the number of observations in each group; values for coding and non-coding regions are identical. **(B)** Relationsh ip between genome size and the percentage of non-coding DNA. **(C)** Relationsh ip between genome size and average intergenic length. **(D)** Relationship between genome size and average intron length. **(E)** Relationship between genome size and gene number. Each point corresponds to a different species. Black lines represent the best-fit regression when all eukaryotic species are pooled. Grey shading indicates the 95% confidence intervals for regression analyses of cyanobacteria and eukaryotes. Lines of best fit for individual taxonomic groups are shown when slopes differed sign ificantly from zero and the number of observations exceeded four. For the percentage of non-cod ing DNA, local regression was used instead of standardized major axis regression.

### Mathematical expression and estimation of selection coefficients of indels

The quota model ^19^ was used to derive the selection coefficients of indel mutations, *s*, from the nutrient cost of indels (Δ*Q*_*min*_) in 83 species with available parameters on cellular minimum quota (*Q*_*min*_) (see methods; figs. S2 and S3, and table S1 ^20,25–28^):

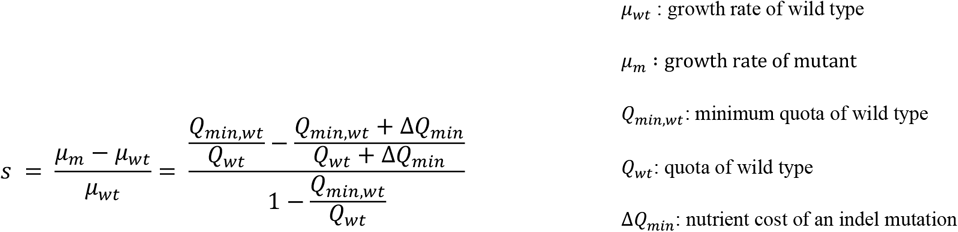

The absolute value of selection coefficients can be derived for the effect of phosphate (*s*_*P*_) and nitrogen (*s*_*N*_) limitation (Supplementary Method section S1). As expected, the selection coefficient increases with the size of the indel, with deletions yielding positive *s* values and insertions yielding negative ones (Fig. 2A and fig. S4). When the contribution of indels to the DNA-packaging N cost of histones is included (*s*_*Nh*_), the relationship with indel size remains monotonic but shows sharp changes whenever a new nucleosome is added or removed. Consequently, the same indel size may correspond to two different *s*_*Nh*_ values, depending on nucleosome addition (or removal) (Fig. 2A). Thus, *s*_*Nh*_ depends not only on indel size but also on chromosomal context, as nucleosome number can vary across loci ^29^.

**Fig. 2.**
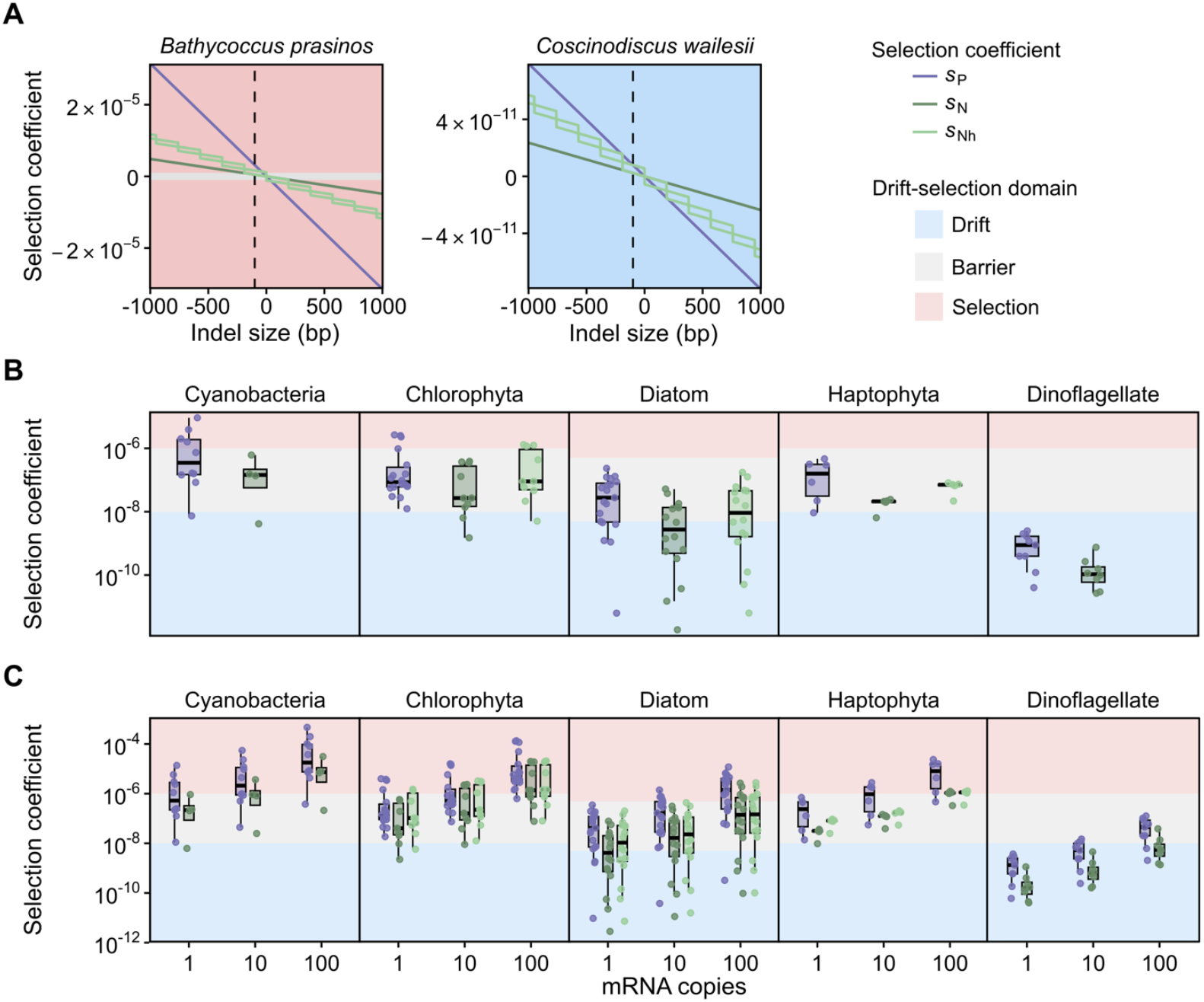
Selection coefficient estimates from the theoretical approach and available quotas and growth rates in phytoplankton. (A) Selection coefficients as a function of indel size in untranscribed regions for the chlorophyte *Bathycoccus prasinos* (low minimum quotas for both P and N: *Q*_minP_ = 0.09 fmol P cell^-1^ and *Q*_minN_ = 2.14 fmol N cell^-1^) and the diatom *Coscinodiscus wailesii* (large *Q*_min_: *Q*_minP_ = 3.50 × 10^4^ fmol P cell^-1^ and *Q*_minN_ = 4.40 × 10^5^ fmol N cell^-1^). Selection coefficients were calculated using Eq. 1 and reflect the impact of indels on P (*s*_P_) or N (*s*_Nh_ and *s*_N_, with or without considering the histone N) requirements under P-or N-limiting conditions. Panel background colors indicate the relative importance of drift (blue) versus natural selection (pink) in indel fixation as indicated in the legend. The vertical dashed line marks a 100-bp deletion, with the corresponding selection coefficients represented in panel B. Plots for 83 individual species, including transcribed regions, are available in Fig. S5. (B) Selection coefficients for 100-bp deletions in untranscribed regions (each point represents one species). Legend as in panel A. A version including species identities is provided in Fig. S5. (C) Selection coefficients for 100-bp deletions in transcribed regions at different expression levels (i.e., number of mRNA copies). Species identities are shown in Fig. S6.

The estimated selection coefficients allow comparison of the fixation probability of a non-transcribed (i.e. number of copies of mRNA, *c*_mRNA_, = 0) 100 bp deletion across different species (Fig. 2A-C and figs. S4 and S5). Selection coefficients *s*_P_, *s*_*N*_, and *s*_*Nh*_ range six orders of magnitude, from 10^-13^ to 10^-7^ (Fig. 2B and fig. S5). Between-species variation in selection coefficients is driven by *Q*_*min*_ which spans more than five orders of magnitude. The minimum *s* value required for selection to be effective depends on the inverse of the effective population size, *N*_e_, such that *N*_e_·s_>_1 (2*N*_e_·s_>_1 in diploids). In phytoplankton, *N*_e_ ranges from 10^6^ to 10^8^ and is typically around 10^7^ (table S2) ^11^. Accordingly, we consider selection effective when *s* > 10^-7^ for haploids and *s* > 5 10^-8^ for diploids. These thresholds are met in 28 out of 70 for *s*_*P*_, in 8 out of 47 for *s*_*N*_, and in 10 out of 35 species for *s*_*Nh*_ (fig. S4). Selection coefficients can be approximated by the fractional cost of an indel, Δ*Q*_*min*_/*Q*_*min*_. In species with low *Q*_*min*_ — such as *Prochlorococcus* and *Mamiellales* — 100 bp indel mutations have selection coefficients such that *N*_e_·s>1 (fig. S4). Selection coefficients may exceed the conservative threshold of 10^-6^ for larger indels or those occurring within introns of highly expressed genes (Fig 2D and fig. S5). In contrast, in species with large *Q*_*min*_ — such as *Coscinodiscus* diatoms or large dinoflagellates — intergenic indels must exceed 1 Mb to reach selection coefficients on the order of |*s*|∼10^-7^. These estimates show that P and N limitation can significantly influence selection on indel mutations in dispensable non-coding regions, which have little or no impact on cellular function, particularly in phytoplankton species with low cellular P and N requirements.

### Stronger selection coefficients associated to P as compared to N limitation

Of note, the selection coefficient associated with phosphorus cellular requirements, *s*_*P*,_ was higher than that for nitrogen, *s*_*N*,_ in all species, and higher than *s*_*Nh*_ in 18 out of 25 species (fig. S5). Overall, phosphate requirements exert a stronger selective pressure on indels than nitrogen requirements. The relative impact of P versus N limitation on the selection of a given indel can be approximated by comparing the Δ*Q*_*min*_/*Q*_*min*_ ratios for P and N. Thus, the relative impact of P versus N limitation will be particularly high in taxa with high N requirements in relative terms, such as cyanobacteria and Chlorophyta, whereas it will be more balanced in groups with low cellular N:P ratios, such as diatoms and Haptophyta ^30^. Indeed, the six species with *s*_*P <*_ *s*_*Nh*_ — five diatoms and two raphidophytes, all diploids — have *Q*_*minN*_/*Q*_*minP*_ <13. In transcribed regions, *s*_*P*_ more frequently exceeds *s*_*Nh*_ (Fig. 2D and figs. S4 and S6), because the number of RNA copies amplifies changes in P (and N) in nucleotides, while histone-associated N remains unaffected. In addition, the *s* values are affected by the actual quota and growing conditions; *s* approaches 0 when *Q* >> *Q*_min_, i.e. under no nutrient limitation and maximal growth (fig. S7). Therefore, for most species, except those with low *Q*_minN_/*Q*_minP_ ratios, environments where P is the primary limiting nutrient exert stronger selection for genome reduction than those limited by N.

### Estimation of the fate of duplications

Gene duplications are a major type of indel mutations (4). To investigate the fate of duplications, we compared the minimum length of indels required for selection to be effective, *l*_min_, to the lengths of non-coding regions. In eukaryotic species with available *N*_e_ estimates, the minimum indel length in non-transcribed regions ranged from 7–1717 bp under phosphorus limitation and 47–2755 bp under nitrogen limitation, with the lowest values observed in *Ostreococcus tauri* (fig. S8). In *Prochlorococcus marinus, l*_min_ was even smaller—only 3 bp under P-limitation. Interestingly, in most species, the estimated *l*_min_ values—calculated assuming *N*_e_ = 10^8^ and *N*_e_ = 10^6^—are similar to the median sizes of intergenic regions (fig. S8). Indeed, the 99th percentile of intergenic region sizes exceeded *l*_min_ in the majority of species. For transcribed regions, *l*_min_ was one-third lower than for non-transcribed regions, and, in most species, also aligned with the median intron size (fig. S8). This alignment between *l*_min_ and the average size of non-coding regions suggests that, on average, duplications larger than the average intergenic region are subject to effective purifying selection, or conversely, that deletions of such duplicated segments may be positively selected.

### Positive correlation between genome size and the minimum cellular nutrient content *Q*_***min***_

A corollary expectation of higher selection coefficients against insertions —or for deletions — in species with low *Q*_*min*_, is a positive correlation between the current size of genomic regions and *Q*_*min*_. Indeed, genome sizes, as well as the sizes of the different genomic regions (non-coding, coding, intergenic and intronic), correlate positively with *Q*_*min*_ measures across eukaryotic species and genera, both for *Q*_*minP*_ and *Q*_*minN*_ (Fig. 3, fig. S9 and table S3). These correlations remain significant after excluding the *Prorocentrum spp*. outlier observation and focusing on species-level data (p-value < 0.05, except for the average intron lengths), and are also significant for the green algae and cyanobacteria subsets (Fig. 3 and fig. S9, tables S4 and S6). To ensure these positive relationships were not soeley driven by the P and N content of DNA, we repeated the analyses using *Q*_*min*_ values corrected by subtracting the N and P content of the relevant genome compartment (fig. S10 and table S5). All correlations remained significant after correction, implying that DNA nitrogen and phosphorus content alone does not fully explain the positive relationship between DNA compartment lengths and *Q*_*min*_.

**Fig. 3.**
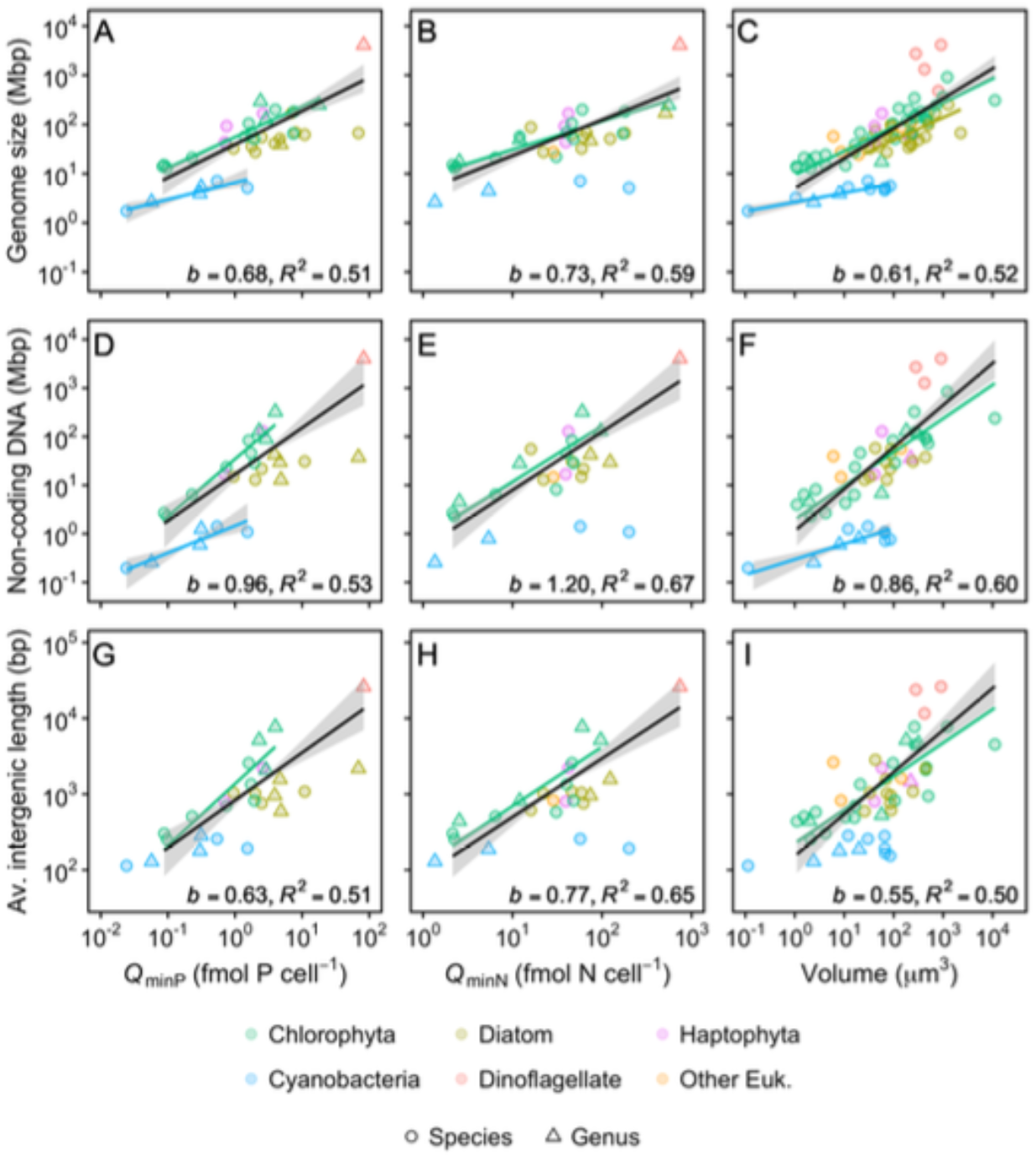
Relationship between genomic traits and *Q*_minP_, *Q*_minN_ and cell volume on a log-log scale. (A, B, C) Genome size. (D, E, F) Total non-coding DNA. (G, H, I) Average size of intergenic regions. Each point represents a different species or genus (see methods). Point color and shape indicate taxonomic affiliation and taxonomic level, respectively. Note that the set of species and the x-axis scales are not the same for *Q*_*minP*_, *Q*_*minN*_, and cell volume. Black lines represent best-fit regressions when all eukaryotic species are pooled, with slope coefficients, *b*, and explained variances, *R*^2^, reported at the bottom of each panel. Grey shading indicates the 95% confidence interval for regressions of cyanobacteria and eukaryotes. Best-fit regressions for individual taxonomic groups are shown when slopes were significantly different from zero and the number of observations exceeded four.

Given the long-standing interest in genome size–cell volume relationships ^5,31–33^, we investigated the influence of cell volume on genomic region size. The slope coefficients relating the size of DNA regions to cell volume were positive and similar to those observed for the relationships between DNA regions and *Q*_min_, particularly *Q*_minP_. This is as expected because cell volume and *Q*_min_ are positively correlated ^20^ (Fig. 3, fig. S9 and table S6). Furthermore, we estimated the variance in genome size explained by *Q*_min_ and cell volume from partial regression analysis (table S7). Most of the explained variation in genome and DNA regions size is shared between *Q*_min_ and cell volume. Thus, our model and these analyses suggest that cellular nutrient requirements, rather than cell volume *per se*, are the primary drivers of the relationship between genome size and cell volume in phytoplankton. Nevertheless, this does not exclude the possibility that selection of smaller cell volumes, that increase the surface to volume ratio and maximizes nutrient uptake in oligotrophic environments^34^, may ultimately affect *Q*_min_ and thus indirectly promote genome reduction.

### Observed intragenomic variations in intron and intergenic region lengths

The model further predicts that selection coefficients increase with transcription rate (Δ*Q*_*min*_ increases with *c*_mRNA_; Eqs. S3-S5). In the low-*Q*_min_ *Ostreococcus* species, selection will be effective for non-transcribed indels longer than 12 bp under P-limiting conditions, which corresponds, in terms of the P budget, to a 1bp deletion with 22 mRNA copies. Consequently, shorter deletions are more likely to become fixed in highly transcribed regions, leading to a negative relationship between the length of introns and their transcription rates. Indeed, introns in the higher transcription-rate class are significantly shorter than those in regions with lower transcription rates in *Bathycoccus* and *Ostreococcus* (Fig. 4 and fig. S12). Compact genomes such as those of *Ostreococcus* and *Bathycoccus* show transcription at up to 99% of intergenic sites, with transcription rates spanning four orders of magnitude (figs. S11 and S12). Intergenic region size decreases with increasing expression, but shows a slight increase in the highest expression class, consistent with the presence of functional elements in these regions (figs. S11-12). The elemental cost of mRNA thus provides an indirect explanation for the origin of intragenomic variation in intron^35^ and, to a lesser extent, in intergenic lengths.

**Fig. 4.**
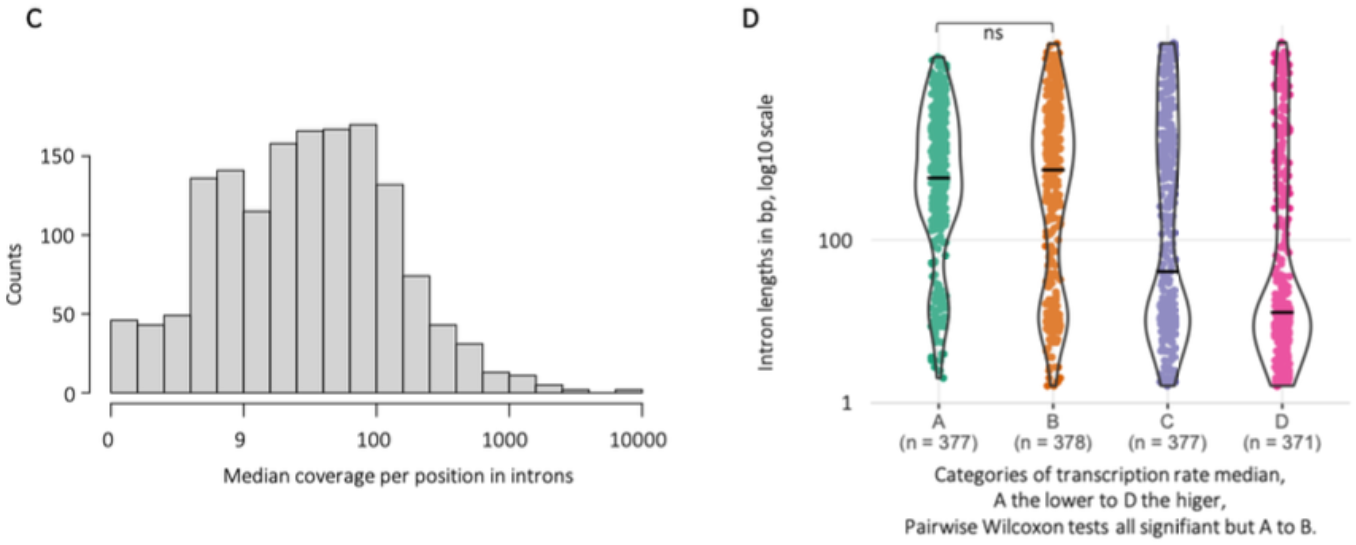
Transcription rates of introns in *Bathycoccus prasinos*. (C) Distribution of median RNA coverage across introns. (D) Median RNA coverage across introns grouped by length category.

### Indirect evidence of selection against insertions in more oligotrophic environments

Selection may also leave signatures on mutations still segregating within natural populations ^36^. Under prevalent limiting nutrient conditions, we expect insertions and deletions to be more and less deleterious, respectively, and all the more so in longer indels. This should be reflected in a lower minor allele frequency (MAF) of insertions as compared to deletions, as well as a lower MAF in longer insertions ^37^. These predictions were tested by leveraging metagenomic and associated environmental data from the TARA Oceans pan-oceanic survey ^38,39^, focusing on species whose presence has been documented at 68 out of 133 sampling stations ^40^. By mapping reads against the reference genomes, we identified 8260 indels with a coverage >= 20 reads, located in intergenic regions, and uniquely present in one metagenome (Fig. 5A, see methods section 9). Of these indels, 4871 (59%) and 2100 (25%) were affiliated to the cosmopolitan *Bathycoccus prasinos* and *Ostreococcus lucimarinus* species (fig. S14). Consistent with our hypothesis, MAF of long insertions were significantly lower than MAF of short insertions in lower P (W = 23069, p < 0.001; Fig. 5B) and lower N (W = 24242, p < 0.05; fig. S13B and table S8) environments. Moreover, the MAF of long insertions (≥10bp) was markedly lower in waters with a lower P concentration than in waters with a higher P concentration (Wilcoxon, W = 91.5, p < 0.01; Fig. 5B). MAF comparisons for long insertions between environments with lower and higher N concentrations were qualitatively similar but non-significant (fig. S13 and table S8). Similar results were obtained when analyses were performed for *B. prasinos* and *O. lucimarinus* separately, or when analyzing median MAF values per metagenome (fig. S14, S15 and table S8). Thus, selection on indels driven by phosphate economy may also be detected on shorter evolutionary timescales, that is, on polymorphisms segregating within natural populations.

**Fig. 5.**
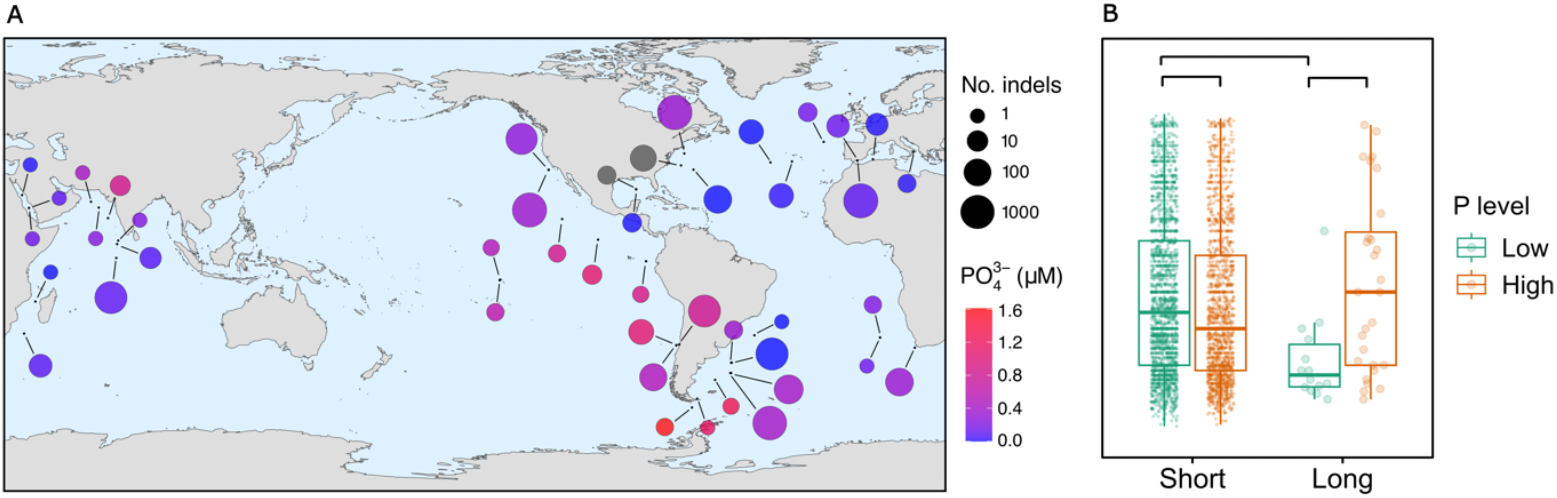
Minor allele frequency (MAF) of insertions and deletions as a function of indel length and environmental phosphate concentration. (A) Worldwide distribution of metagenomes containing polymorphism for the target species. (B) MAF of short (<10bp) and long (>=10bp) indels in low (green) and high (orange) phosphorus environments. Results of the pairwise comparisons corresponding to panel B are provided in Table S8.

## Conclusions

In conclusion, we provide an empirically informed theoretical background of the processes underlying genome streamlining observed in phytoplankton inhabiting oligotrophic waters (e.g. *Prochlorococcus* and *Pelagibacter*) ^1,7,16,41^. Previous pan-oceanic metagenome analyses have reported shorter genome sizes in prokaryotes from tropical than polar oceans, which was mainly attributed to temperature effects ^42^. Our alternative explanation highlights the role of nutrient limitation and the higher selective pressures to promote genome compactness of photoautotrophic organisms in oligotrophic subtropical waters.

Unlike heterotrophs, phytoplankton rarely experience carbon limitation, and its potential effect on genome size is likely lower than that of nitrogen (N) and phosphorus (P), given cellular and nucleotide stoichiometry ^43^. Energy limitation, i.e. light in photoautotrophs, can certainly contribute to genome reduction, and light limiting conditions often alternate with nutrient limitation in oceans and lakes ^10,44^. However, selective pressures exerted by N and especially P limitation may be stronger. To evaluate this, one may compare the main determinants of selection coefficients for indels under nutrient limitation (Δ*Q*_min_/*Q*_min_) versus energy cost (Δ*E*/*E:* cell energy budget) as reported in Lynch and Marinov (2015)^10^. Note that this comparison is relevant for the replication associated costs of non-coding regions. We found that, in eukaryotic species Δ100 μm^3^, the fractional costs in terms of P and N were one order of magnitude and nearly five times higher, respectively, than energy costs. This corresponds to Δ*Q*_minP_/*Q*_minP_ Δ 2 Δ*Q*_minN_/*Q*_minN_ ≈ 10 ΔE/E (fig. S16). This is particularly relevant under the current global change scenario, which might enhance P limitation due to anthropogenic N releases and a more marked stratification ^45–48^.

The model suggests several evolutionary implications of N and P constraints on genome evolution. First, higher *s* values in species with low *Q*_min_ reflect the higher relative costs of nucleotides in terms of N and P, which implies that each decrease in *Q*_min_ further strengthens selection for deletions. Second, any variation in *Q*_min_, which can be promoted by a given indel or by any other cellular change, affect the strength of nutrient-mediated selective pressures on future mutations. This generates positive feedback on genome size: a large insertion, such as a whole chromosome duplication, alters the selection efficiency against subsequent indels. Third, changes in *Q*_min_ can result from multiple small indels at different loci, not necessarily a single event, and thus permit effective selection against insertion-biased mutator alleles. Fourth, relaxed selection in high *Q*_min_ species increases the likelihood that duplications or horizontally transferred sequences are retained compared with low *Q*_min_ species. Thus, neofunctionalization and genome diversification following gene duplication are limited in low *Q*_min_ species or ecotypes thriving in oligotrophic environments, but more likely in nutrient-rich environments.

## Material and methods are provided as supplementary material

## Acknowledgments

We are grateful to all Genophy group members for stimulating discussions and comments, especially Frederic Sanchez, Anais Labecot and Julien Ferrero for help with the cultures and pilot experimentations. We would like to thank the GenoToul bioinformatic platform for access to the computing facilities and David Pecqueur and Christophe Salmeron the BIOPIC platform, for access and support to the cytometry facilities.

## Author contributions

Conceptualization: CC, GP

Methodology: CC, MK, OC, SG, GP

Investigation: CC, MK, GP

Funding acquisition: CC, SG, GP

Writing – original draft: CC, GP

Writing – review & editing: CC, MK, SG, GP

## Data and materials availability

All data are available in the main text or the supplementary materials.

## Supplementary Materials Materials and Methods Supplementary Text

Figs. S1 to S16 Tables S1 to S8

References(*1*–*45*)

Data S1 to S3

## References

1. Giovannoni, S.J., Tripp, H.J., Givan, S., Podar, M., Vergin, K.L., Baptista, D., Bibbs, L., Eads, J., Richardson, T.H., Noordewier, M., et al. (2005). Genetics: Genome streamlining in a cosmopolitan oceanic bacterium. Science (1979) 309, 1242–1245. 10.1126/science.1114057.

2. Fernández, P., Amice, R., Bruy, D., Christenhusz, M.J.M., Leitch, I.J., Leitch, A.L., Pokorny, L., Hidalgo, O., and Pellicer, J. (2024). A 160 Gbp fork fern genome shatters size record for eukaryotes. iScience 27. 10.1016/j.isci.2024.109889.

3. Petrov, D.A. (2001). Evolution of genome size: new approaches to an old problem. TRENDS in Genetics 17, 23–28.

4. Lynch, M., and Conery, J.S. (2003). The Origins of Genome Complexity.

5. Gregory, T.R. (2001). Coincidence, coevolution, or causation? DNA content, cellsize, and the C-value enigma. Biological Reviews 76, 65–101. 10.1111/j.1469-185X.2000.tb00059.x.

6. Swan, B.K., Tupper, B., Sczyrba, A., Lauro, F.M., Martinez-Garcia, M., Gonźalez, J.M., Luo, H., Wright, J.J., Landry, Z.C., Hanson, N.W., et al. (2013). Prevalent genome streamlining and latitudinal divergence of planktonic bacteria in the surface ocean. Proc Natl Acad Sci U S A 110, 11463–11468. 10.1073/pnas.1304246110.

7. Giovannoni, S.J., Cameron Thrash, J., and Temperton, B. (2014). Implications of streamlining theory for microbial ecology. Preprint at Nature Publishing Group, 10.1038/ismej.2014.60 https://doi.org/10.1038/ismej.2014.60.

8. Lauro, F.M., Mcdougald, D., Thomas, T., Williams, T.J., Egan, S., Rice, S., Demaere, M.Z., Ting, L., Ertan, H., Johnson, J., et al. (2009). The genomic basis of trophic strategy in marine bacteria 10.1073/pnas.0903507106.

9. Okie, J.G., Poret-Peterson, A.T., Lee, Z.M.P., Richter, A., Alcaraz, L.D., Eguiarte, L.E., Siefert, J.L., Souza, V., Dupont, C.L., and Elser, J.J. (2020). Genomic adaptations in information processing underpin trophic strategy in a whole-ecosystem nutrient enrichment experiment. Elife 9. 10.7554/eLife.49816.

10. Lynch, M., and Marinov, G.K. (2015). The bioenergetic costs of a gene. Proc Natl Acad Sci U S A 112, 15690–15695. 10.1073/pnas.1514974112.

11. Lynch, M.R. (2024). Evolutionary cell biology: the origins of cellular architecture (Oxford University Press).

12. Martiny, A.C., Coleman, M.L., and Chisholm, S.W. (2006). Phosphate acquisition genes in Prochlorococcus ecotypes: Evidence for genome-wide adaptation.

13. Ustick, L.J., Larkin, A.A., Garcia, C.A., Garcia, N.S., Brock, M.L., Lee, J.A., Wiseman, N.A., Moore, J.K., and Martiny, A.C. (2021). Metagenomic analysis reveals global-scale patterns of ocean nutrient limitation. Science (1979) 372, 287–291.

14. Lin, S., Litaker, R.W., and Sunda, W.G. (2016). Phosphorus physiological ecology and molecular mechanisms in marine phytoplankton. J Phycol 52, 10–36.

15. Moore, C.M., Mills, M.M., Arrigo, K.R., Berman-Frank, I., Bopp, L., Boyd, P.W., Galbraith, E.D., Geider, R.J., Guieu, C., Jaccard, S.L., et al. (2013). Processes and patterns of oceanic nutrient limitation. Nat Geosci 6, 701–710.

16. Dufresne, A., Garczarek, L., and Partensky, F. (2005). Accelerated evolution associated with genome reduction in a free-living prokaryote. Genome Biol 6, 1–10.

17. Derelle, E., Ferraz, C., Rombauts, S., Rouzé, P., Worden, A.Z., Robbens, S., Dé Ric Partensky, F., Degroeve, S., Echeynié, S., Cooke, R., et al. (2006). Genome analysis of the smallest free-living eukaryote Ostreococcus tauri unveils many unique features.

18. Palenik, B., Grimwood, J., Aerts, A., Rouzé, P., Salamov, A., Putnam, N., Dupont, C., Jorgensen, R., Derelle, E., Rombauts, S., et al. (2007). The tiny eukaryote Ostreococcus provides genomic insights into the paradox of plankton speciation.

19. Droop, M.R. (1968). Vitamin B12 and marine ecology. IV. The kinetics of uptake, growth and inhibition in Monochrysis lutheri. Journal of the Marine Biological Association of the United Kingdom 48, 689–733.

20. Edwards, K.F., Thomas, M.K., Klausmeier, C.A., and Litchman, E. (2012). Allometric scaling and taxonomic variation in nutrient utilization traits and maximum growth rate of phytoplankton. Limnol Oceanogr 57, 554–566. 10.4319/lo.2012.57.2.0554.

21. Sibbald, S.J., and Archibald, J.M. (2020). Genomic insights into plastid evolution. Genome Biol Evol 12, 978–990. 10.1093/GBE/EVAA096.

22. Gregory, T.R. (2005). The C-value enigma in plants and animals: A review of parallels and an appeal for partnership. In Annals of Botany, pp. 133–146. 10.1093/aob/mci009.

23. Michael, T.P. (2014). Plant genome size variation: Bloating and purging DNA. Brief Funct Genomic Proteomic 13, 308–317. 10.1093/bfgp/elu005.

24. Pellicer, J., Hidalgo, O., Dodsworth, S., and Leitch, I.J. (2018). Genome size diversity and its impact on the evolution of land plants. Preprint at MDPI AG, 10.3390/genes9020088 https://doi.org/10.3390/genes9020088.

25. Ebenezer, V., Hu, Y., Carnicer, O., Irwin, A.J., Follows, M.J., and Finkel, Z. V. (2022). Elemental and macromolecular composition of the marine Chloropicophyceae, a major group of oceanic photosynthetic picoeukaryotes. Limnol Oceanogr 67, 540–551. 10.1002/lno.12013.

26. Liefer, J.D., Garg, A., Fyfe, M.H., Irwin, A.J., Benner, I., Brown, C.M., Follows, M.J., Omta, A.W., and Finkel, Z. V. (2019). The macromolecular basis of phytoplankton C:N:P under nitrogen starvation. Front Microbiol 10. 10.3389/fmicb.2019.00763.

27. Marañón, E., Cermeño, P., López-Sandoval, D.C., Rodríguez-Ramos, T., Sobrino, C., Huete-Ortega, M., Blanco, J.M., and Rodríguez, J. (2013). Unimodal size scaling of phytoplankton growth and the size dependence of nutrient uptake and use. Ecol Lett 16, 371–379. 10.1111/ele.12052.

28. Chevin, L.M. (2011). On measuring selection in experimental evolution. Biol Lett 7, 210– 213. 10.1098/rsbl.2010.0580.

29. Almassalha, L.M., Carignano, M., Pujadas Liwag, E., Shun Li, W., Gong, R., Acosta, N., Dunton, C.L., Carrillo Gonzalez, P., Carter, L.M., Kakkaramadam, R., et al. (2025). Chromatin conformation, gene transcription, and nucleosome remodeling as an emergent system.

30. Ho, T.Y., Quigg, A., Finkel, Z. V., Milligan, A.J., Wyman, K., Falkowski, P.G., and Morel, F.M.M. (2003). The elemental composition of some marine phytoplankton. J Phycol 39, 1145–1159. 10.1111/j.0022-3646.2003.03-090.x.

31. Cavalier-Smith, T. (2005). Economy, speed and size matter: Evolutionary forces driving nuclear genome miniaturization and expansion. In Annals of Botany, pp. 147–175. 10.1093/aob/mci010.

32. Connolly, J.A., Oliver, M.J., Beaulieu, J.M., Knight, C.A., Tomanek, L., and Moline, M.A. (2008). Correlated evolution of genome size and cell volume in diatoms (Bacillariophyceae). J Phycol 44, 124–131. 10.1111/j.1529-8817.2007.00452.x.

33. Wang, H., Wu, P., Xiong, L., Kim, H.S., Kim, J.H., and Ki, J.S. (2024). Nuclear genome of dinoflagellates: Size variation and insights into evolutionary mechanisms. Preprint at Elsevier GmbH, 10.1016/j.ejop.2024.126061 https://doi.org/10.1016/j.ejop.2024.126061.

34. Litchman, E., Klausmeier, C.A., Schofield, O.M., and Falkowski, P.G. (2007). The role of functional traits and trade-offs in structuring phytoplankton communities: Scaling from cellular to ecosystem level. Ecol Lett 10, 1170–1181. 10.1111/j.1461-0248.2007.01117.x.

35. Castillo-Davis, C.I., Mekhedov, S.L., Hartl, D.L., Koonin, E. V., and Kondrashov, F.A. (2002). Selection for short introns in highly expressed genes. Nat Genet 31. 10.1038/ng940.

36. Shenhav, L., and Zeevi, D. (2020). Resource conservation manifests in the genetic code.

37. Sawyer, S.A., and Hartlt, D.L. (1992). Population Genetics of Polymorphism and Divergence.

38. Sommeria-Klein, G., Watteaux, R., Ibarbalz, F.M., Karlusich, J.J.P., Iudicone, D., Bowler, C., and Morlon, H. (2021). Global drivers of eukaryotic plankton biogeography in the sunlit ocean. Science (1979) 374, 594–599. 10.1126/science.abb3717.

39. Sunagawa, S., Acinas, S.G., Bork, P., Bowler, C., Babin, M., Boss, E., Cochrane, G., de Vargas, C., Follows, M., Gorsky, G., et al. (2020). Tara Oceans: towards global ocean ecosystems biology. Preprint at Nature Research, 10.1038/s41579-020-0364-5 https://doi.org/10.1038/s41579-020-0364-5.

40. Leconte, J., Benites, L.F., Vannier, T., Wincker, P., Piganeau, G., and Jaillon, O. (2020). Genome resolved biogeography of mamiellales. Genes (Basel) 11. 10.3390/genes11010066.

41. Partensky, F., and Garczarek, L. (2010). Prochlorococcus: Advantages and limits of minimalism. Ann Rev Mar Sci 2, 305–331. 10.1146/annurev-marine-120308-081034.

42. Ngugi, D.K., Acinas, S.G., Sánchez, P., Gasol, J.M., Agusti, S., Karl, D.M., and Duarte, C.M. (2023). Abiotic selection of microbial genome size in the global ocean. Nat Commun 14. 10.1038/s41467-023-36988-x.

43. Sterner, R.W., and Elser, J.J. (2002). Ecological Stoichiometry: The Biology of Elements from Molecules to the Biosphere (Princeton University Press).

44. Longhurst, A. (2007). Ecological geography of the sea (Academic Press).

45. Karl, D.M. (2014). Microbially mediated transformations of phosphorus in the sea: New views of an old cycle. Ann Rev Mar Sci 6, 279–337. 10.1146/annurev-marine-010213-135046.

46. Burson, A., Stomp, M., Akil, L., Brussaard, C.P.D., and Huisman, J. (2016). Unbalanced reduction of nutrient loads has created an offshore gradient from phosphorus to nitrogen limitation in the North Sea. Limnol Oceanogr 61, 869–888. 10.1002/lno.10257.

47. Duhamel, S., Diaz, J.M., Adams, J.C., Djaoudi, K., Steck, V., and Waggoner, E.M. (2021). Phosphorus as an integral component of global marine biogeochemistry. Preprint at Nature Research, 10.1038/s41561-021-00755-8 https://doi.org/10.1038/s41561-021-00755-8.

48. Gerace, S.D., Yu, J., Moore, J.K., and Martiny, A.C. (2025). Observed declines in upper ocean phosphate-to-nitrate availability. Proc Natl Acad Sci U S A 122. 10.1073/pnas.2411835122.

